# Evidence for a multicellular symplasmic water pumping mechanism across vascular plant roots

**DOI:** 10.1101/2021.04.19.439789

**Authors:** Valentin Couvreur, Adrien Heymans, Guillaume Lobet, Xavier Draye

## Abstract

With global warming, climate zones are projected to shift poleward, and the frequency and intensity of droughts to increase, driving threats to crop production and ecosystems. Plant hydraulic traits play major roles in coping with such droughts, and process-based plant hydraulics (water flowing along **decreasing pressure Ψ_p_ or total water potential Ψ_tot_ gradients**) has newly been implemented in land surface models.

An enigma reported for the past 35 years is the observation of water flowing along **increasing water potential gradients** across roots. By combining the most advanced modelling tool from the emerging field of plant micro-hydrology with pioneering cell solute mapping data, we found that **the current paradigm of water flow across roots of *all vascular plants* is incomplete: it lacks the impact of solute concentration (and thus negative osmotic potential Ψ_o_) gradients across <u>living</u> cells**. This gradient acts as a water pump as it reduces water tension without loading solutes in plant vasculature (xylem). Importantly, water tension adjustments in roots may have large impacts in leaves due to the tension-cavitation feedback along stems.

Here, we mathematically demonstrate the water pumping mechanism by solving water flow equations analytically on a triple-cell system. Then we show that the simplistic upscaled equations hold in 2- and 3-D maize, grapevine and Arabidopsis complex hydraulic anatomies, and that water may flow “uphill” of water potential gradients toward xylem as observed experimentally.

Besides its contribution to the fundamental understanding of plant water relations, this study lays new foundations for future multidisciplinary research encompassing plant physiology and ecohydrology, and has the ambition to mathematically capture a keystone process for the accurate forecasting of plant water status in crop models and LSMs.

**Graphical Abstract:** 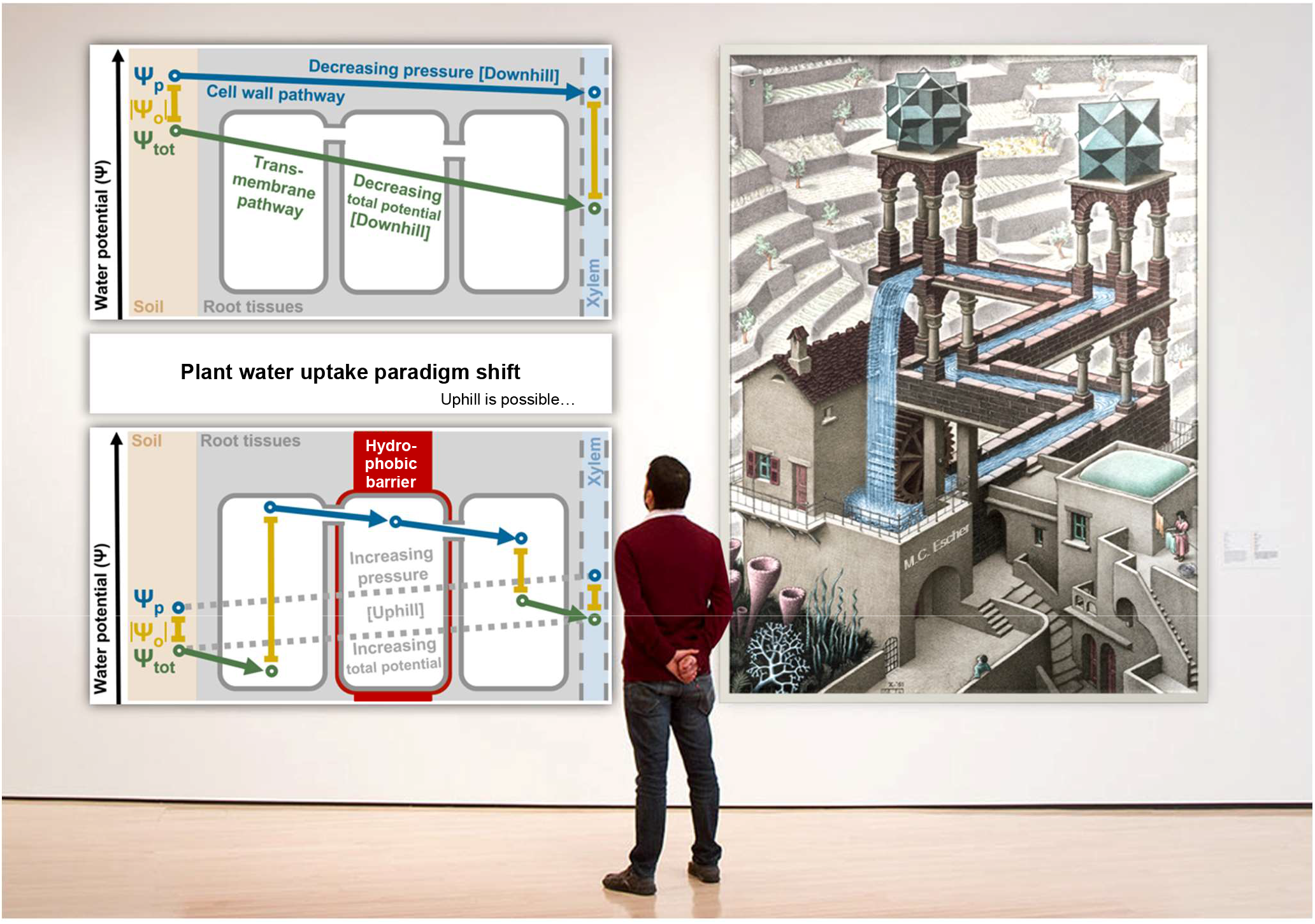

**Highlights:**

- We provide a scale-consistent solution of water flow equations across root tissues
- Symplasmic osmotic potential gradients are missing in the current theory of root water uptake
- The model solves the empirical enigma of root water uptake uphill of water potential gradients

## Introduction

Water flow inside plant tissues is analogous to river streamflow. As rivers flow from high to low elevations, the plant transpiration stream flows along a cascade of water potentials – or free energy status – that are high (in soil) to low (in roots, then even lower in leaves); a theory untouched for more than seven decades (Van Den Honert, 1948). Metaphorically, water flows “downhill” of gradients of summed pressure and gravitational potentials across non-selective tissues, such as xylem vessels, the part of vascular tissues made of dead cells emptied from their protoplasts. That is why cohesive water columns in xylem passively flow against the gravitational pull, toward leaf mesophyll cell walls under extremely negative pressure relative to free water (Steudle, 2001). In contrast, cell cytosol pressure potential is generally positive, thereby allowing cell turgidity and expansion. Such adjustment of water pressure in plants would not be possible without one key component: the selective permeability of cell membranes for major solutes (Kramer and Boyer, 1995). The affinity of water for solutes favours its spontaneous movement across cell membranes toward the side with highest solute concentration. This process called osmosis adds a third component to the potential of water: the osmotic potential (more negative at higher solute concentration). Hence, water flows downhill of gradients of this “total” water potential Ψ_tot_ (sum of gravitational, pressure Ψ_p_ and osmotic potentials Ψ_o_) across membranes decoupling the transport of water and solutes. Membranes thereby allow the coexistence of highly concentrated environments under high pressure (cell cytosols and organelles forming the “symplast”, white areas in Fig. 1) at the same total potential as more diluted environments under lower – often negative – pressure (cell walls and xylem, forming the “apoplast”, light grey and blue areas in Fig. 1, respectively).

**Figure 1.**
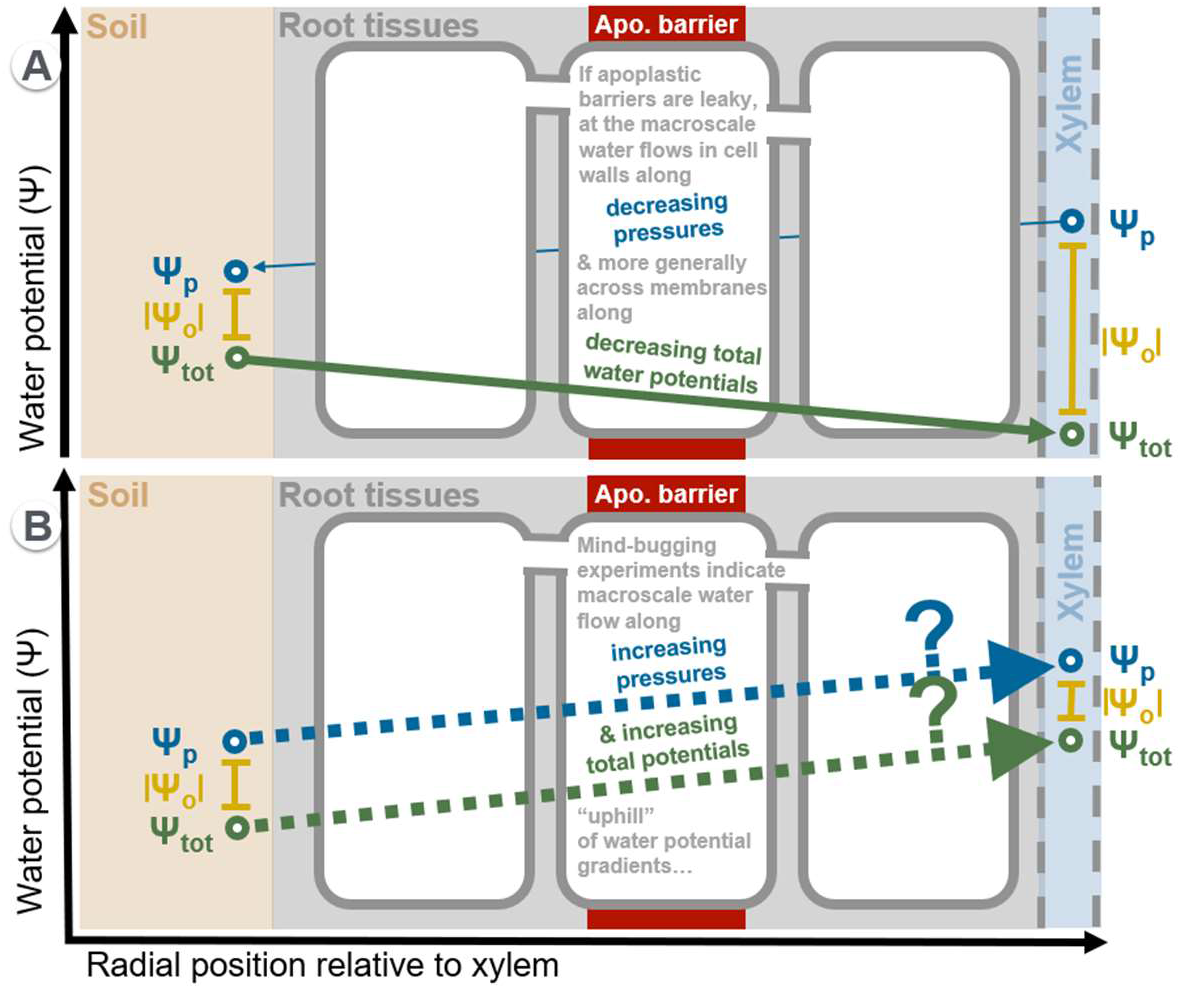
(A) Current paradigm of root radial water flow here in case of solute loading in xylem vessels, and (B) scheme of contradictory experimental results reported in the literature.

It took a masterpiece of natural engineering, the “Casparian strip”, for plant roots to elevate the relative pressure of an entire apoplastic compartment. Its principle: forming a tight scaffold limiting water and solute permeation across radial walls of the endodermal cell layer (Doblas et al., 2017), making it possible to actively accumulate solutes on the xylem-side of the scaffold. With a higher solute concentration, this compartment’s total water potential is lowered (Fig. 1A, right side), inducing water flow across endodermal membranes, into xylem vessels, downhill of the total potential gradient (thick green arrow). The newly found equilibrium with higher xylem pressure may possibly generate pressure-driven water back-flow across cell walls if the Casparian strip is leaky (thin blue arrow bypassing membranes in Fig. 1A).

Water as a solvent may theoretically drag solutes across a leaky fraction of a membrane. When that is the case, the osmotic potential difference across the membrane is not equally effective at moving water, and the driving force is corrected by a multiplicative factor σ called “reflection coefficient” (Katchalsky and Curran, 1967) (equal to one in case of fully selective path, down to zero for paths fully permeable to solutes, e.g. primary cell walls and membranous sleeves – called plasmodesmata – connecting neighbouring cells cytosols). The current paradigm of water flow from root surface to xylem in vascular plants simply conceptualizes root tissues as a “big-membrane” (Kramer and Boyer, 1995) characterised by (i) a radial hydraulic conductivity Lp_r_ [L P^−1^ T^−1^] and (ii) a dimensionless reflection coefficient σ_r_ transposed to the root scale and typically close to unity (Knipfer and Fricke, 2010):

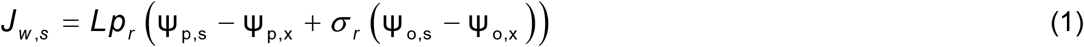

where J_w,s_ [L T^−1^] is water flux at root surface, Ψ_p,s_ and Ψ_p,x_ [P] are the water pressure potentials at root surface and in xylem, respectively, and Ψ_o,s_ and Ψ_o,x_ [P] the (negative) osmotic potentials at root surface and in xylem, respectively.

As active transmembrane water pumps have not been discovered (water flows passively across channel proteins called aquaporins, AQP), the current paradigm of root radial water flow concludes that water only flows “downhill” of Ψ gradients between root surface and xylem. However, numerous studies have reported that plant roots may absorb water “uphill” of both Ψ_p_ and Ψ_tot_ gradients (Fig. 1B, e.g. (Miller, 1985; Enns et al., 2000; Rowan et al., 2000; Bai et al., 2007)). These paradoxical observations have remained a curiosity for the past 35 years, and suggest plants may take up more water from their environment than predicted by theory (Singh, 2016). Small steps toward solving this case have been taken in recent decades. Firstly, the passive water transport across transmembrane channel proteins (aquaporins) could be complemented with direct water pumping by a hypothetical water-ion cotransporter as found in animal epithelial cells (Wegner, 2014). Secondly, Pickard (2003) showed that a hypothetical plasmalemma supporting two parallel water fluxes (one driven by total potential gradient, and the other by pressure potential gradient) would do the trick. Finally, with a microscale computational model of root hydrodynamics, Couvreur et al. (2018a) demonstrated mathematically that radial flow rates deviate from the current radial water flow theory if symplasmic osmotic potentials are non-uniform. Strikingly, water flow “uphill” of gradients of all components of the water potential, from root surface to xylem, was predicted by the hydraulic anatomy model when using an osmotic potential distribution measured by EDX-microanalysis in maize roots (Enns et al., 2000), suggesting all elements needed to solve the “riddle of uphill water flow” were standing under our very eyes.

The importance of this mechanism stands in the discovery of how plants may (i) regulate their root water transport in a way complementary to aquaporin regulation (Chaumont et al., 2005), (ii) increase their water extraction from soils, and (iii) reduce xylem water tension without relying on xylem solute loading (mostly limited to −0.3 MPa under water deficit (Enns et al., 2000; Westhoff et al., 2008)) or stomatal closure (Gleason et al., 2019). Xylem water tension regulation is of particular importance as extreme values lead to catastrophic cavitation in crops (McCully, 1999) and trees, which operate with a narrow hydraulic safety margin (Choat et al., 2012). Cavitation magnifies frictions, which intensify xylem water tension, provoking additional cavitation in a sensitive positive feedback loop compounded along stems (Couvreur et al., 2018b). Thus, root symplasmic osmotic gradients reducing xylem water tension by a fraction of MPa may leverage relatively large effects on canopy water potential and supply, by limiting upstream cavitation in the plant hydraulic continuum.

The objectives of this study are to (i) provide a multiscale solution of radial water flow in a simplistic root hydraulic network as a cell triplet, (ii) demonstrate that the solution applies to realistic root hydraulic anatomies, and (iii) quantify the impact of cell scale osmotic potential distributions observed experimentally on the driving force of water flow across roots. With these steps completed, a new scale-consistent paradigm of root radial water flow will be proposed, ready to be challenged against data from cutting-edge techniques of root osmotic mapping (Persson et al., 2016) and soil-plant water status monitoring (Jerszurki et al., 2017).

## Results

### A multiscale solution of water flow equations across a simple root hydraulic network

In order to produce an upscaled solution of water flow equations in root hydraulic anatomies, we first developed an analytical solution for a simplistic system of three cells. Figure 2 displays composite radial water pathways as hydraulic conductances connecting nodes spanning compartments from the root surface to the xylem. We call pathways associated to flow rates Q_1_ and Q_4_ [L^3^ T^−1^] “cell-to-cell” (i.e. through cell membranes and plasmo-desmata)(Steudle and Peterson, 1998), and those of Q_2_ and Q_3_ “apoplastic” (i.e. through cell walls, except across the endodermis, where membranes are crossed). For simplicity, here we assume no “purely apoplastic” radial water pathway through both primary cell walls and Casparian strip, as excluded by Knipfer and Fricke (2010). The state-of-the-art water flow equations through cell walls, membranes, and plasmodesmata from Kramer and Boyer (1995) for the triple cell system are solved in the methods section. In order to condense the writing of the analytical solution, clusters of cell-scale hydraulic parameters are given names and symbols at the root segment scale (detailed expressions of root segment-scale parameters as functions of cell-scale parameters in the methods section). The analytical solution of the microscale equations, hereafter referred to as “upscaled model”, then yield the following form of the net radial water flux at root surface, J_w,s_ [L T^−1^] (here ratio of Q_1_ + Q_2_ or of Q_3_ + Q_4_ to root surface area, A_s_ [L^2^]):

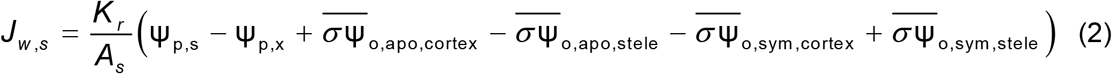

where *K*_r_ [L^3^ T^−1^ P^−1^] is the conductance of the hydraulic network (detailed expression in the methods section, Eq. (27)), Ψ_p,i_ [P] is the pressure potential at the i-th node (note that subscripts “s” and “x” replace the node number at root surface and xylem nodes), and the average osmotic potentials in the symplast and apoplast (subscripts “apo” and “sym”) on the cortex and stele sides originate from the following expressions:

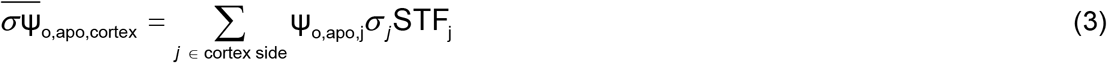

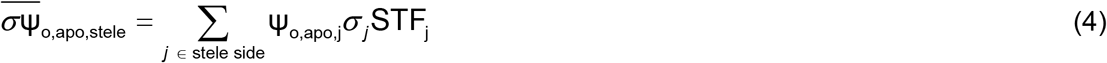

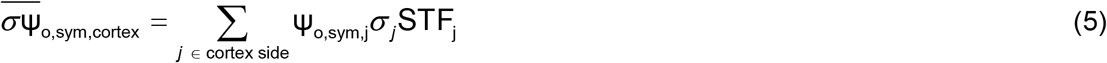

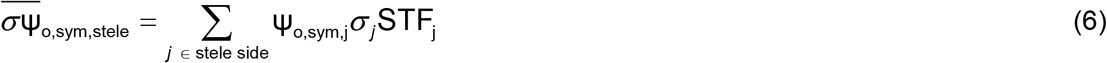

where *j* is the membrane number (note that opposed endodermal membranes have different numbers, hence the symbols K_m2_ and K_m3_, where the former belongs to the cortex side, while the latter belongs to the stele side), σ_*j*_ [dimensionless] is the solute reflection coefficient of the membrane, Ψ_o,apo,*j*_ and Ψ_o,sym,*j*_ [P] are the osmotic potentials on the apoplastic and symplasmic sides of the membrane, respectively, and STF_*j*_ [dimensionless] is the membrane “Standard Transmembrane Fraction” (detailed expression in the methods section, Eqs. (24-25)) which equals the fraction of water flowing through membrane *j* in conditions of uniform osmotic potential (i.e. “standard” condition). The latter property implies that STF_*j*_ values integrate to 1 on both the cortex- and the stele-sides, so that the left-hand-sides of Eqs. (3-6) can be considered as weighted-average osmotic potentials (corrected by reflection coefficients), hence the “bar” symbol on top.

**Figure 2.**
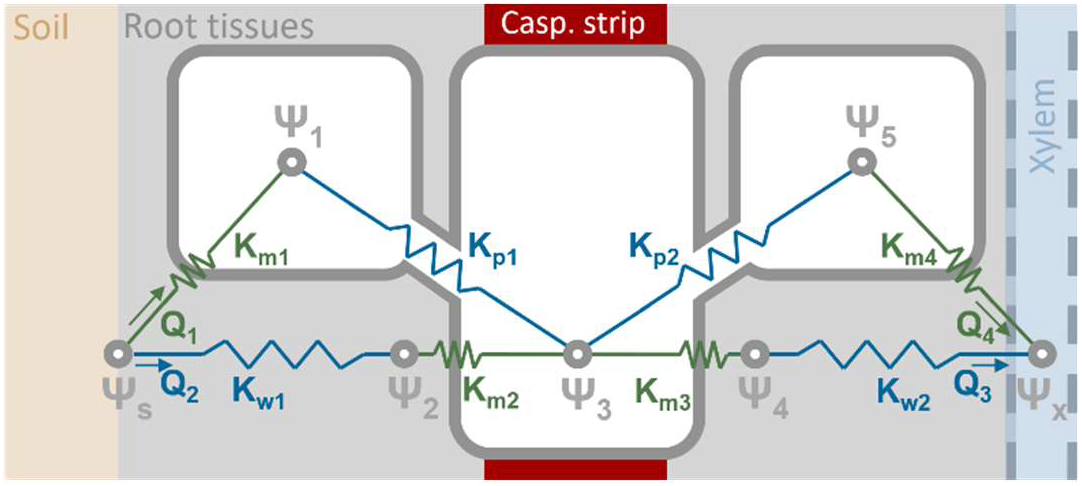
Scheme of the simplistic hydraulic network used to derive an analytical solution of water flow along root water paths. The three cells have dimensions modified to ease reading. Ψ_p_ and Ψ_tot_ gradients drive water flow along blue and green paths, respectively.

The solution of J_w,s_ expressed in Eq. (2) is multiscale as it emerges from cell-scale water flow equations, while only properties and variables of the system as a whole eventually remain. The upscaled expression shares similarities and a couple striking differences with the independent mainstream model (Eq. 1) of radial water flow across roots segments (Kramer and Boyer, 1995), which will be discussed further down.

### The multiscale solution of water flow holds in diverse complex root hydraulic anatomies

A comparison of microscale simulations of water fluxes at root surface (J_w,s_) with both mainstream and multiscale macroscopic models predictions is provided in Fig. 3, in *Zea mays* (maize), *Vitis riparia* (grapevine) and *Arabidopsis thaliana* primary roots. Maize and *A. thaliana* anatomies were digitised from microscopic images, using the software CellSet (Pound et al., 2012), while the grapevine anatomy including clusters of aerenchyma was generated with the root cross-section simulator GRANAR (Heymans et al., 2020). Water fluxes were computed in networks of cell walls, membranes and plasmodesmata using the micro-hydrological model MECHA (Couvreur et al., 2018a). Axially, simulations cover the thickness of one cell length in maize and grapevine, so that the geometry is essentially two-dimensional, while it is three-dimensional in *A. thaliana*, with 250 layers connected axially cell-to-cell and along cell walls (Fig. 3A-C, note that only a fraction of fluxes in cell walls is displayed to ease visualisation). Due to the dominant apoplastic water transport, water fluxes in cell wall tends to increase from the outer to the inner layers of the cortex, before passing through the symplast in the vicinity of the endodermis, and reaching early metaxylem vessels. Water fluxes in cell walls also increase substantially in areas of convergence, such as septa separating aerenchyma spaces (Fig. 3B). Higher water fluxes in *A. thaliana* distal cell walls (left side in Fig. 3C) are due to the suberisation of endodermal walls in the proximal region, limiting radial water flow, while the distal side only bears a Casparian strip.

**Figure 3.**
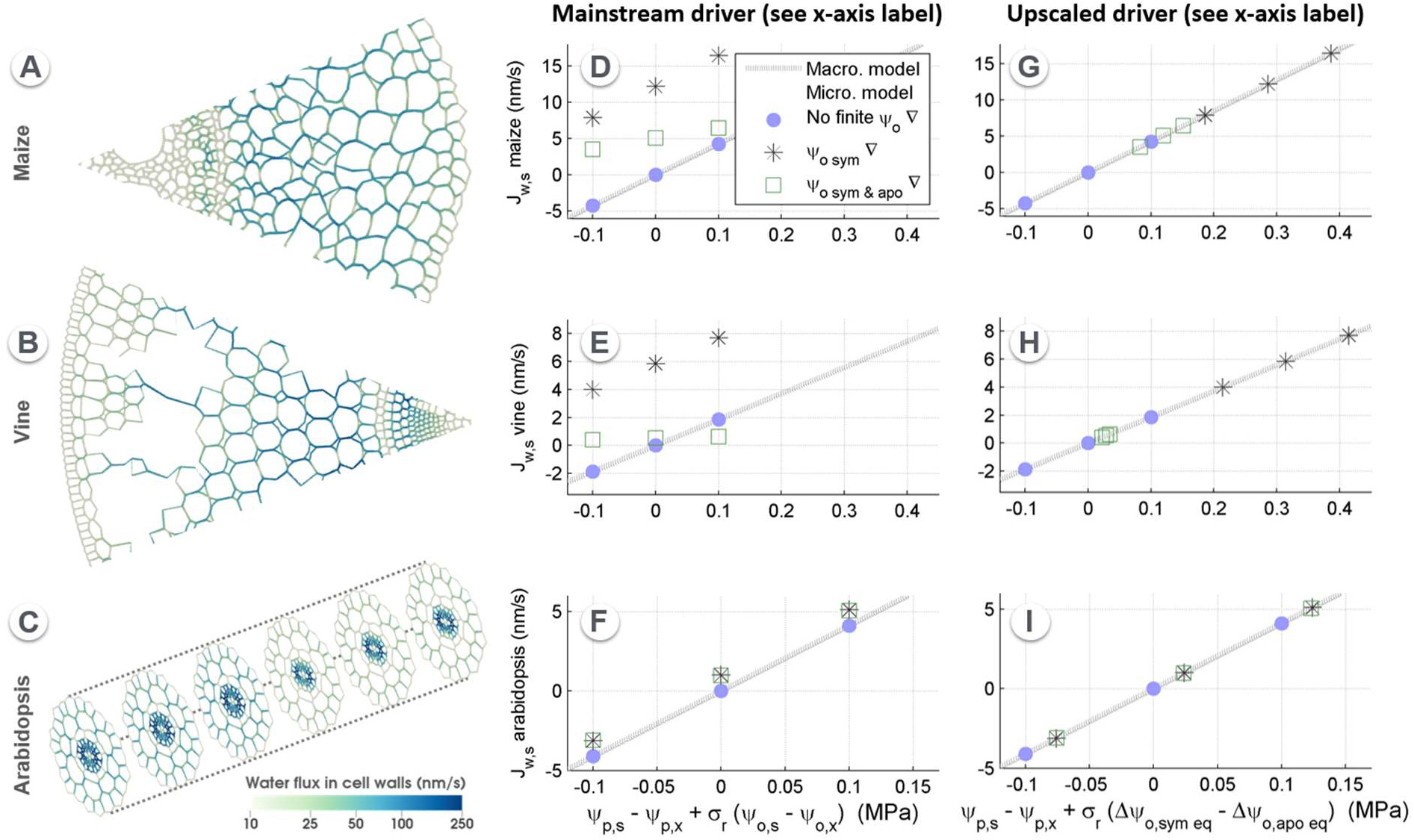
Comparison of water fluxes in 3 models of root water uptake: microscale, mainstream macroscale, and upscaled macroscale. (A-C) Partial view of microscale water fluxes simulated with MECHA in maize, vine, and *A. thaliana*. (D-F) Water fluxes at root surface simulated with MECHA (J_w,s_) are not proportional to driving forces of the mainstream macroscale radial water flow model, except in absence of osmotic gradients within the symplast and within the apoplast. (G-I) Driving forces of the proposed upscaled macroscale model of radial water flow align with J_w,s_ simulated from the microscale.

When setting uniform osmotic potentials (Ψ_o_) within the symplast, as well as within the apoplastic compartments on the stele- and cortex-sides of the endodermis, J_w,s_ values simulated from the microscale (disks in Fig. 3D-F) align with the mainstream macroscopic model (grey line, Eq. 1). However, microscale predictions strongly deviate from the mainstream macroscale model when Ψ_o_ is non-uniform within the symplast (stars in Fig. 3D-F, using gradients reported in the literature in maize (Rygol et al., 1993; Enns et al., 2000)), or both within the symplast and apoplastic compartments (squares in Fig. 3D-F, microscale apoplastic Ψ_o_ gradients then being simulated with a model of solute convection-diffusion (Couvreur et al., 2018a)). Note that each of the aforementioned osmotic scenarios is simulated with the microscale model under high, medium, and low xylem water pressure (Ψ_p,x_ of 0.02, −0.08, and −0.18 MPa, respectively), hence the tripling of each symbol. Based on these triplets, apparent root radial hydraulic conductivities (*Lp*_r,app_) in the scenario with both types of osmotic gradients (squares in Fig. 3) were calculated as the slope of the J_w,s_ response to variations of Ψ_p,x_. Ratios of *Lp*_r,app_ to *Lp*_r_ differed across plants. In vine, *Lp*_r,app_ was only 7% of *Lp*_r_, while it was 36% in maize, and was left almost unaltered in *A. thaliana* (99%).

Overall the correlation coefficients R^2^ for the microscale J_w,s_ versus driving force of mainstream macroscopic model were 0.22, 0.13, and 0.98, in maize, vine, and *A. thaliana*, respectively in the selected scenarios. Strikingly, the upscaled macroscopic model derived from the simple hydraulic network (Fig. 2) remains a fully exact solution of microscale water flow equations for the complex hydraulic anatomies (R^2^ = 1.00, Fig. 3G-I), including a 3-D root. Importantly, the upscaled parameters *K_r_* and STF neither depend on pressure nor on osmotic potentials within the system or at its boundaries. Therefore, for each root type, parameter values were calculated by solving water flow equations in a single “homogeneous” scenario with null Ψ_o_, null Ψ_p_ at root surface, and non-null Ψ_p_ on the proximal side of xylem vessels. This single parametrization was then used in all 9 scenarios per plant in Fig. 3G-I.

## Discussion

### Current and novel paradigms of radial water flow across roots

In this study we present a novel equation of water flow across root segments (Eq. 2) based on an analytical solution of water flow in a simplistic root composite hydraulic network (Fig. 2). Its mathematical development was a twist in search for a general analytical solution of water flow equations in complex root hydraulic anatomies. This procedure mimics the one applied to produce a simple macroscopic root water uptake model based on the hydraulic architecture approach (Couvreur et al., 2012), in which a mathematical solution of xylem water potential at plant collar in a two-branch root was found to remain exactly valid in complex root systems. Here we demonstrate that the upscaled equation (from sub-cellular to root segment scale) holds in the complex root hydraulic anatomies of maize, grapevine, and *A. thaliana* (Fig. 3, accommodating 2-/3-D geometries, aerenchyma, and non-uniform osmotic potentials in cell walls and protoplasts). The proposed equation (Eq. 2) is in line with the original work of Van Den Honert (1948) on water flow in plants as a “catenary process” (i.e. limited by the most resistive path in series, like in a supply chain), further developed by Kramer and Boyer (1995) who presented the root segment as an “osmometer” (i.e. able to build up water pressure by osmosis after accumulating solutes in xylem vessels). While the current paradigms of water flow at the cell scale (e.g. Eqs. 8-10 in the methods section) and root segment scale (Eq. 1) are well accepted by the scientific community (Kramer and Boyer, 1995), our multiscale analysis shows that they do not necessarily comply with each other, and calls for a revision of the model of the “root osmometer” (Eq. 1), which seems incomplete. In particular, it is neither sensitive to osmotic potential gradients within the apoplast, which may explain biased estimations of the apparent “osmotic” root *Lp*_r_ (Knipfer and Steudle, 2008; Couvreur et al., 2018a), nor to osmotic potential gradients within the symplast observed experimentally (Rygol et al., 1993; Enns et al., 2000), which may offer a solution to the riddle of the apparent “uphill” root water flow (Miller, 1985; Rowan et al., 2000; Pickard, 2003; Bai et al., 2007; Singh, 2016).

As compared to our generalized upscaled model (Eq. 2), the current paradigm of root radial water flow (Eq. 1) appears to be a specific case, which assumes that (i) the average product of osmotic potentials by membrane reflection coefficient on both sides of the endodermis are equal 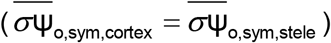, (ii) the average product of osmotic potentials by membrane reflection coefficient in cell walls on the cortex side equal the product of the osmotic potential at root surface by the root reflection coefficient 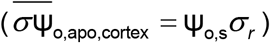, and (iii) the average product of osmotic potentials by membrane reflection coefficient in cell walls on the stele side equal the product of the osmotic potential in xylem by the root reflection coefficient 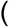 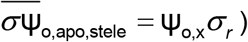. Based on symplasmic osmotic potentials reported by Enns et al. (2000) in maize, the error margin due to the first assumption would be in the range of 0.15 to 0.3 MPa depending on the degree of suberisation of the root endodermis. The error margins of the second and third assumptions essentially depend on the magnitude of radial water fluxes (as diffusion tends to even out solutes when the water convective component is low), osmotic potentials at root surface and in xylem (e.g. there would be no substantial accumulation of solute in the vicinity of the endodermis if solute concentration at root surface was negligible), and the length of the path along which solutes may accumulate (e.g. particularly long in grapevine and short in *A. thaliana*). Our results suggest that this quantitative error margin may range between 0.0 MPa (here in *A. thaliana*) and 0.4 MPa (here in grapevine under root xylem water pressure of −0.18 MPa). Such errors may in part cancel out, or add up, depending on their respective signs.

It is worth noting that Zhu and Steudle (1991) also proposed a multiscale solution of water flow between root segment and cell scales to estimate cell hydraulic properties from root scale measurements. Their top-down approach accounts for the areas, hydraulic conductivities, and reflection coefficients of cell walls and protoplasts. It also leaves the door open to bidirectional parallel water fluxes through the cell-to-cell and purely apoplastic pathways as our model does, while neglecting osmotic potential gradients within the root and pressure-driven flow through plasmodesmata.

### Deciphering the symplasmic mechanism of water pumping

An illustration of the concept of radial water flow across roots emerging from multiscale hydraulic principles is summarized in Fig. 4B-C. The symplasmic Ψ_o_ gradient (yellow bars of decreasing size from the cortex to the pericycle, from about −0.8 to −0.5 MPa in Enns et al. (2000) control conditions), tend to generate a gradient of cell turgidity (i.e. water pressure). As plasmodesmata are not selective for major osmolites, water flow is essentially pressure-driven from the cortex under high pressure to the pericyle under lower pressure (blue arrows pointing to the right). Interestingly, on this symplasmic path, water flows uphill of the Ψ_tot_ gradient (which does not come into play as no selective membrane is crossed), so that Ψ_tot_ on the symplasmic side of the pericycle membrane may be larger than on the symplasmic side of the cortex membrane. Water naturally flows downhill of Ψ_tot_ gradients across these membranes, completing the pathway from root surface to xylem. Note that in Fig. 4, arrow thickness stands for flow rate, and that endodermis suberization prevents any backflow of water in the 3^rd^ panel, making this version of the water pump more efficient. Besides, high solute concentration in phloem sieve tubes does not generate a substantial outward driving force for water, but only bi-directional water flow with its neighbouring cells, due to the absence of apoplastic barrier separating them.

**Figure 4.**
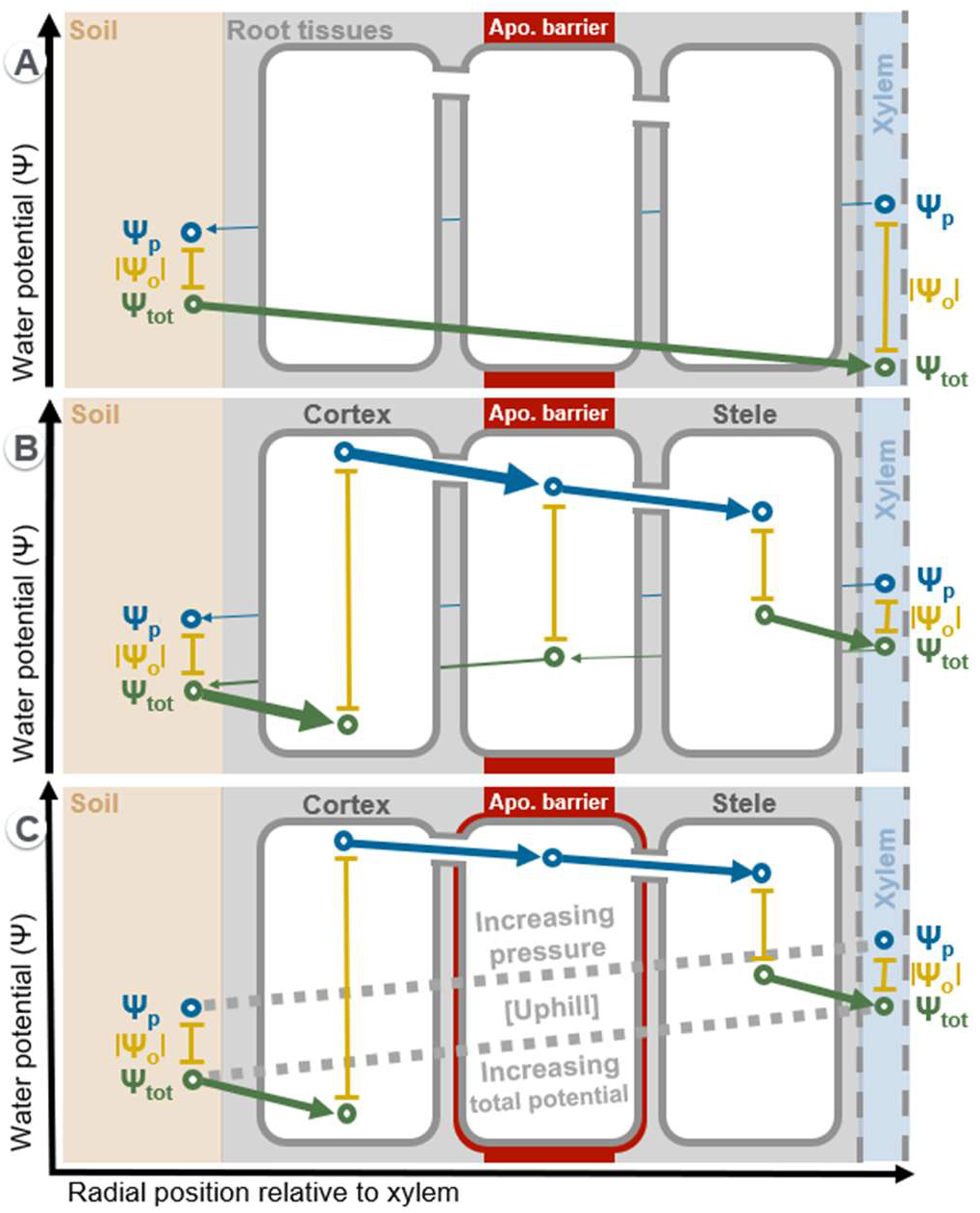
Mainstream “apoplastic” (A) and emerging “symplasmic” (B, C) concepts of root water indirect pumping.

Because Ψ_o_ gradients in the apoplast (as in Fig. 4A) and in the symplast (as in Fig. 4B-C) may both contribute to indirect water pumping, we suggest to distinguish them as apoplastic and symplasmic water pumping mechanisms. They may be combined, and rely on the maintenance of Ψ_o_ gradients, which requires energy for transmembrane solute transfer or solute metabolism.

### Outlook

Like in a magic trick orchestrated by vascular plants, the attention of the scientific community has been focalised on a part of the driving force of water uptake (water potentials at root surface and in xylem). When water kept flowing “uphill” of macro-scale water potential gradients, the fascinated researcher conjectured on the existence of an active transmembrane water pump (Wegner, 2014), or even questioned the applicability of the second law of thermodynamics in plants (Bai et al., 2007). Our mathematical results combined to experimental observations by Enns et al. (2000) support that the trick may lie in a “concealed” osmotic gradient along living cells, driving water flow in an impossible direction according to the current paradigm of water uptake by plant roots. It is particularly mind-bending that at the micro-scale water may systematically flow downhill of the relevant components of water potential (Ψ_p_ or Ψ_tot_, depending on the local path) while simultaneously flowing uphill of both Ψ_p_ and Ψ_tot_ gradients at the root cylinder macro-scale. Such a tale is reminiscent of the universe of M.C. Escher, in which all may seem to flow downhill from close range, though the surprise comes as we take a step back (see Graphical Abstract).

Beyond the proposition of a mathematical solution to the “uphill water flow enigma” reported experimentally, our work calls for an in-depth documentation of osmotic potential distribution in living root cells, now facilitated by the cutting-edge technology of LA-ICP-MS (Persson et al., 2016), in order to capture the dynamics of this concealed driving force of water uptake. In the meantime, simplifying assumptions need to be done, as it has always been the case in mainstream root water uptake models. The proposed emergent model allows dropping some of the most limiting assumptions of mainstream models, such as the absence of osmotic potentials in living cells, and absent plasmodesmatal pathway in equations of radial flow (virtually included in the cell-to-cell pathway, but actually neglected at the stage of translation into mathematical equations in Zhu and Steudle (1991)).

The question remains open of how symplasmic osmotic gradients are formed and possibly maintained at a homeostatic state, despite symplasmic solute convection from cells under higher pressure toward their neighbours. Interestingly, Enns et al. (2000) showed that adding substantial amounts of KNO_3_ to the root bathing solution does alter absolute values of osmotic potential distribution in the symplast, but not their gradients across root tissues. Yet, the data available on such solute transfer and metabolism properties is currently insufficient for a complete detailed modelling. However, Couvreur et al. (2018a) (Supplemental Note S2) evaluated that maintaining osmotic potential gradients observed in the symplast of maize roots by Enns et al. (2000) would require solute transport rates lower than typically observed for calcium and nitrate (2.8 10^−7^ mol m^−2^s^−1^ and 6 10^−8^ mol m^−2^s^−1^, respectively) (Cárdenas et al., 1999; Pouliquin et al., 2000). Furthermore, several other processes may contribute to keep osmotic potentials stable, such as the metabolism of large molecules, and other ion transporters. At this stage though, we are left with the option of considering that snapshots of measured solute distributions can be used to simulate snapshots of water flow across roots.

## Methods

In this section, we detail (i) the development of the analytical solution of water flow in the simple hydraulic network, (ii) how macroscopic parameters numerical values can be calculated, and (iii) how the multiscale model is validated against numerical solutions from the microscale hydraulics model MECHA.

### Development of the analytical solution of water flow equations in the simple root hydraulic network

The simple root hydraulic network with associated symbols for components of water potentials “Ψ”, flow rates “*Q*”, and hydraulic conductances of membranes “*K*_m_”, cell walls “*K*_w_” and plasmodesmata “*K*_p_” numbered as in Fig. 2 translate into the following water flow equations:

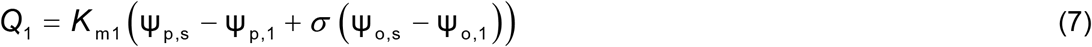

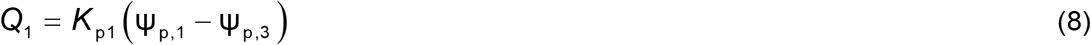

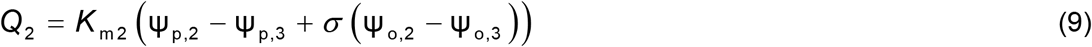

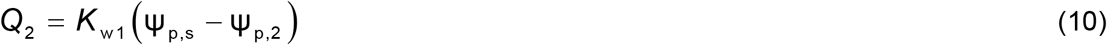

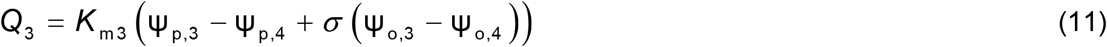

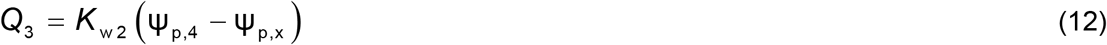

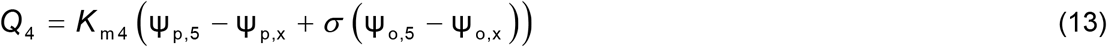

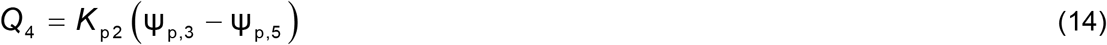

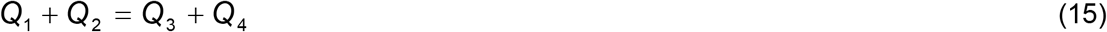

where Ψ_p,i_ and Ψ_o,i_ are the pressure and osmotic potentials at the i-th node, respectively. Note that labels “s” and “x” replacing the node number apply to root surface and xylem boundary nodes, respectively.

Under Dirichlet boundary conditions (prescribed Ψ_p,s_ and Ψ_p,x_), the system has 9 unknowns (Q_1-4_, Ψ_p,1-5_), while the following parameters are considered as “known” variables, which will however keep their non-numerical form in this exercise: *K*_m1-4_, *K*_p1-2_, *K*_w1-2_, Ψ_p,s_, Ψ_o,s_, Ψ_p,x_, Ψ_o,x_, Ψ_o,1-5_, σ_1-4_.

From the membrane numeration point of view in this simple scheme, we also name osmotic potentials on the apoplast and symplast sides of the j-th membrane Ψ_o,apo,j_ and Ψ_o,sym,j_, respectively. Thus, for instance Ψ_o,apo,1_ and Ψ_o,sym,1_ correspond to Ψ_o,s_ and Ψ_o,1_ in Fig. 2, respectively.

In order to simplify the mathematical expression, the equivalent conductance of each path (made of two conductances in series) is referred to as *K*_Qj_ (with j the path number), so that:

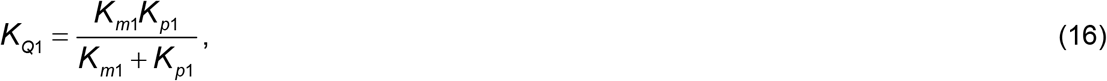

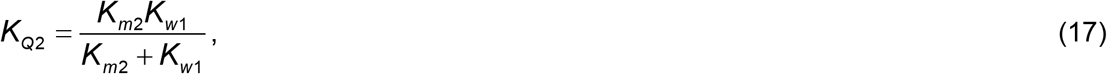

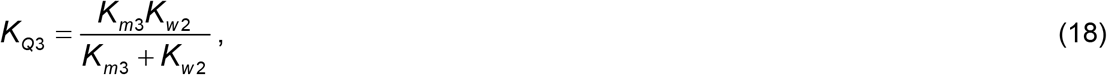

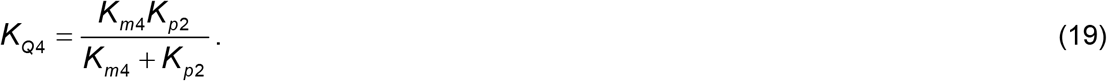

The system of equations 7-15 has the following analytical solution of water flow rate on the cortical side (paths 1 and 2, with j the path number):

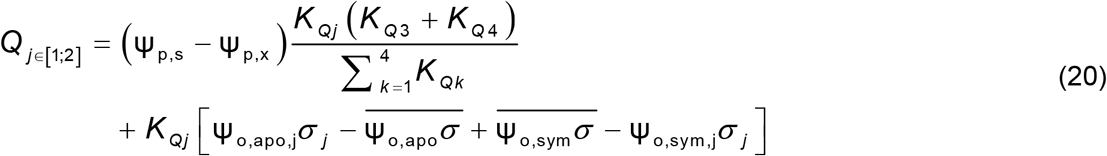

where

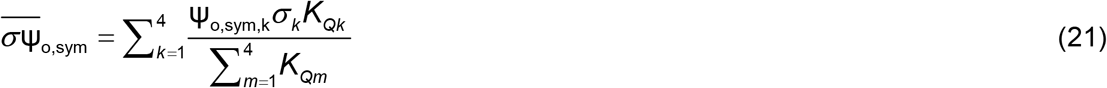

and

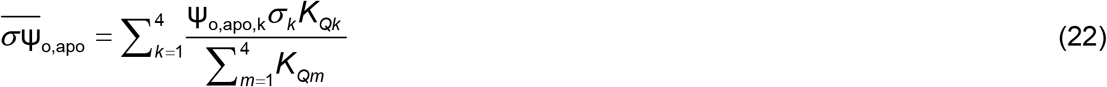

On the stele side (paths 3 and 4, with j the path number), the analytical solution of water flow rate is as follows:

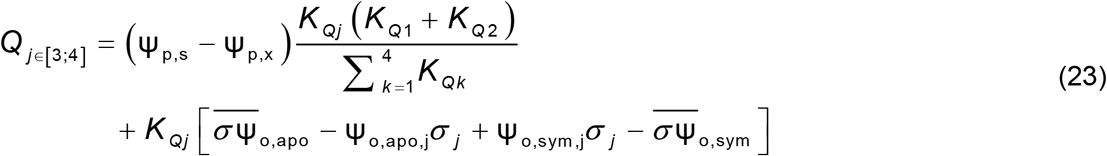

Assuming uniform membrane reflection coefficients, in conditions of negligible gradients of osmotic potential within the apoplast and within the symplast and (as commonly assumed in root water uptake models (Doussan et al., 1998; Bouda et al., 2018)), the osmotic terms cancel out in Eqs. (20-23). Under such conditions, referred to as “standard” in the following, the fraction of the total water flow passing through the transmembrane path “j” on the cortex side (i.e.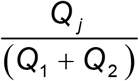) simply corresponds to the following “Standard Transmembrane Fraction” (STF, dimensionless):

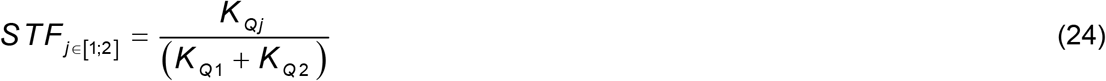

In complement, we have:

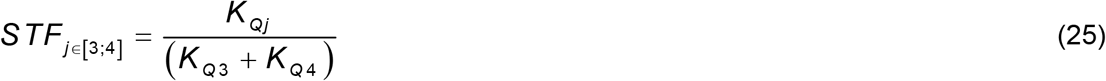

It is then trivial that the integration of Standard Transmembrane Fractions on the cortex side, and on the stele side, both equal 1.

In “standard” conditions (no osmotic gradients within the apoplast or the symplast), the total water flux at root surface (J_w,s_ [m s^−1^], ratio of Q_1_ + Q_2_ or of Q_3_ + Q_4_ to root surface area, A_s_ [m^2^]) simplifies down to:

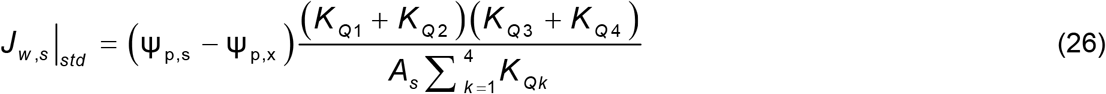

Where the equivalent conductance of the full hydraulic network (*K*_r_ [m^3^ MPa^−1^ s^−1^]) can be isolated:

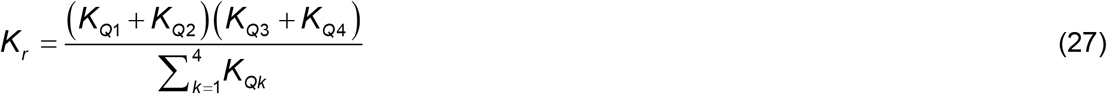

In the following, K_r_ and the STF vector are called “macroscopic parameters” as they aggregate multiple cell scale parameters. They can be used to simplify the writing of the general expression of J_w,s_, this time accounting for osmotic gradients:

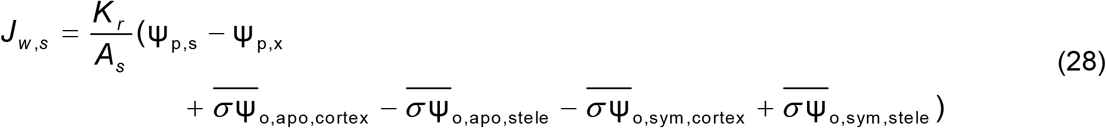

where separate weighted-average values of the osmotic potential are evaluated on the cortex and stele sides for both apoplastic and symplasmic compartments:

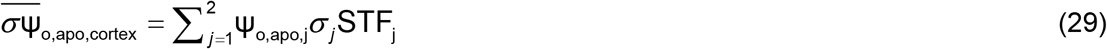

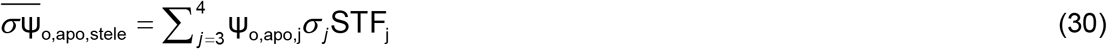

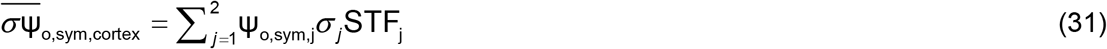

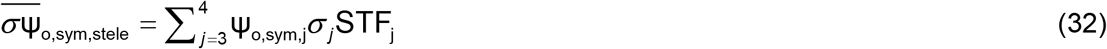

## Notes

### Competing Interest Statement

The authors have declared no competing interest.

